# Finding ancestry-specific chromatin architecture in Alzheimer’s disease

**DOI:** 10.1101/2025.08.19.671054

**Authors:** Wanying Xu, Liyong Wang, Leina Lu, Xiaoxiao Liu, Farid Rajabli, Katrina Celis, Marla Gearing, David A. Bennett, Margaret Flanagan, Sandra Weintraub, Changiz Geula, Theresa Schuck, William K. Scott, Derek M. Dykxhoorn, Anthony J. Griswold, Margaret A. Pericak-Vance, Juan I. Young, Fulai Jin, Jeffery M. Vance

**Affiliations:** Department of Genetics and Genome Sciences, School of Medicine, Case Western Reserve University, Cleveland, OH 44106, USA; The Biomedical Sciences Training Program (BSTP), School of Medicine, Case Western Reserve University, Cleveland, OH 44106, USA; John P. Hussman Institute for Human Genomics, University of Miami Miller School of Medicine, Miami, Florida, FL 33136, USA; John T. Macdonald Foundation Department of Human Genetics, University of Miami Miller School of Medicine, Miami, Florida, FL 33136, USA; Goizueta Alzheimer’s Disease Research Center, Emory University, Atlanta, GA, 30322, USA; Department of Neurological Sciences, Rush University, Chicago, IL, USA; Northwestern ADC Neuropathology Core, Northwestern University Feinberg School of Medicine, Chicago, IL, 60611, USA; The Department of Pathology and Laboratory Medicine, Institute on Aging and Center for Neurodegenerative Disease Research, The Perelman School of Medicine at the University of Pennsylvania, Philadelphia, PA, USA; Department of Computer and Data Sciences, Department of Population and Quantitative Health Sciences, Case Comprehensive Cancer Center, Case Western Reserve University, Cleveland, OH 44106, USA

## Abstract

Genetic risk for Alzheimer’s Disease (AD) varies across populations. We hypothesized that three-dimensional (3D) genome architecture variations could offer novel epigenetic understanding of ancestry-specific genetic risk. Herein, we performed Hi-C analyses of frontal cortex from *APOE* ε4/ε4 individuals with African (AF) or European (EU) ancestry who also had single nuclei ATAC-seq and RNA-seq data available. Ancestry-specific 3D genome architecture was found at both compartment and chromatin loop levels. EU genomes have more active compartments than AF genomes, consistent with our previous report of higher chromatin accessibility in EU than AF genome. Of the over one million chromatin loops identified by the *DeepLoop* pipeline, we called 12,082 putative EU-specific and 2885 putative AF- specific loops. The AF-specific loops are smaller (median size =158 kb) and likely represent promoter-enhancer interactions, while EU-specific loops are larger (median size = 496 kb) and enriched for CTCF loops. We found that differently expressed genes between AF and EU ancestries were significantly enriched at the putative ancestry-specific loop loci (Fisher-test; p<2.2×10^-16^; OR=5.13). High confidence HiC-QTLs (N=38) were identified after filtering with CTCF consensus sequence and chromatin accessibility-QTLs. Our study demonstrates variations in 3D genome structure between ancestries, which may contribute to the ancestry-specific genetic risk.

## Introduction

Most of the variants identified in Genome Wide Association Studies (GWAS) are in non-coding genomic regions, including those for Alzheimer Disease (AD^1^). Current understanding suggests that these disease-associated SNPs are likely to be functional in non-coding cis-regulatory elements (CREs), *e.g.*, enhancer, repressor, or transcription factor binding sites. While the roles of the associated SNPs can be inferred from analyses of co-localization with CRE and quantitative trait loci (QTLs), correlating SNP genotypes with epigenetic traits, such as DNA methylation, DNase I hypersensitivity, chromatin accessibility, and gene expression^2–12^, unraveling the actual genes and mechanisms driving these associations is still a challenge. This is because 1) regulatory elements identified by GWAS can be far from their actual target gene, 2) one regulatory element can target multiple genes, and 3) multiple regulatory elements can target one gene.

Further complicating the assignment of GWAS risk variants to target genes are potential differences in regulatory architecture between ancestries. GWAS in AD have identified over 75 loci in individuals of European (EU) ancestry^10^. Recent GWAS in other populations have demonstrated that genetic architectures for AD vary across different ancestry backgrounds ^13^. One of the best examples of this is the disease risk for Alzheimer’s disease (AD) driven by the presence of the Apolipoprotein 4 allele (*APOE ε4*). The *APOE* ε4 allele is the major genetic risk factor for AD across different populations but with a different effect size between populations.

The strongest effect is in East Asians, followed by Europeans, with the lowest effect in individuals with African (AF) ancestry^10, 14^. We and others have shown that the local genomic ancestry surrounding *APOE* is associated with the different risks between populations ^15–18^. Indeed, European carriers of *APOE* ε4 have greater expression of the gene than African ancestry carriers and have more open chromatin genome wide by ATAC^19, 20^. In addition to the different effect size across populations, ancestry-specific variants and loci have been identified, such as an ATP binding cassette subfamily A member 7 (*ABCA7)* deletion in people with African ancestry^13, 21^. Not surprisingly, the AD polygenic risk score (PRS) derived from European populations perform poorly in other admixed populations, such as African American and Caribbean Hispanics^22^. Understanding the mechanisms underlying the ancestry-specific genetic risk is important for precision medicine and may also provide novel therapeutic opportunities.

The spatial arrangement of DNA in three-dimensional (3D) space influences the accessibility of promoters to regulatory elements such as enhancers and repressors. As such, variations in the 3D genome organization represent a potential epigenetic mechanism for disease risk via perturbation of gene expression. 3D genome structure analysis facilitates target gene identification by providing physical evidence connecting regulatory elements and their target genes, even when they are far away from each other. The Chromosome Conformation Capture (3C) technique has been developed to capture the 3D genome structure. Hi-C is the extension of the 3C technique coupled with next generation sequencing, which enables global mapping of 3D chromatin structure^23, 24^. At a lower-resolution (500 kb) level, the genome can be partitioned into compartment A (transcriptionally active euchromatin) and compartment B (transcriptionally silenced heterochromatin)^25^.. On the other hand, chromatin loops are high-resolution 3D genome features that often represent promoter-enhancer interactions and are believed to play a direct role on gene expression regulation and therefore cell fate decision^5^. Identifying and quantifying chromatin loops from Hi-C data, however, has been challenging due to bias and high noise levels at kilobase resolution^6^.

To overcome this hurdle in the study of 3D genome structure, we recently developed *DeepLoop*^7, 8^, a deep-learning based tool to improve the sensitivity of Hi-C loop analysis. Importantly, we demonstrated that *DeepLoop* improves the robustness of loop strength quantification across different Hi-C platforms. To explore 3D genome structure as an epigenetic mechanism underlying the differential genetic risk for AD across populations, we performed Hi-C analysis in frontal cortex from donors with Alzheimer Disease: four with high global EU ancestry and four with high global AF ancestry. We used *DeepLoop* to quantify chromatin loops and to determine the correlation of genetic variants with loop strength (HiC-QTLs). Our hypothesis is that ancestry-specific genetic variants can contribute to ancestry differences in genetic risk for AD by affecting 3D chromatin interactions. We focused our analysis on ancestry-specific chromatin loops to maximize the effect of genetic variation on chromatin loops, increasing the power to map these HiC-QTLs.

## Results

### Samples

We generated Hi-C libraries from postmortem frontal cortex tissues (Brodmann area 9) from four African American (AA) and four Non-Hispanic White (NHW) donors with Alzheimer’s Disease. All donors are homozygous *for APOE* ε4. All NHW and AA donors are homozygous for EU and AF local ancestry surrounding the *APOE* gene, respectively. These samples have been previously described, with single nuclei ATAC and single nuclei RNAseq data available ^19, 20^. Each of the NHW donors has over 95% EU ancestry and the AA donors have admixed ancestry with an average of 82% AF ancestry. The age-of-death ranged from 71 to 93 years, and the samples are sex matched across the ancestries. Previous studies demonstrated similar cellular proportions^19, 20^ and that European local ancestry is randomly distributed in our AA samples, so any Hi-C data in a particular location most likely reflects African ancestry ^19^.

### Compartment variations between AF and EU ancestry

The Hi-C libraries of each sample were sequenced to an average depth of ∼700M reads (paired-end 150bp), or ∼70X genome coverage after removing PCR duplicates **(Supplementary Table 1)**. The genome was partitioned into compartment A (associated with transcriptionally active euchromatin) and compartment B (associated with transcriptionally silenced heterochromatin) at 500-kb resolution. The state of compartmentalization was compared between EU and AF ancestry. Out of 6,206 compartment bins across the genome, 525 bins showed suggestive evidence for compartment difference (*P* ≤ 0.1; two-side t-test): 499 more active in EU and 26 more active in AF (**Fig. 1b**).

**Figure 1.**
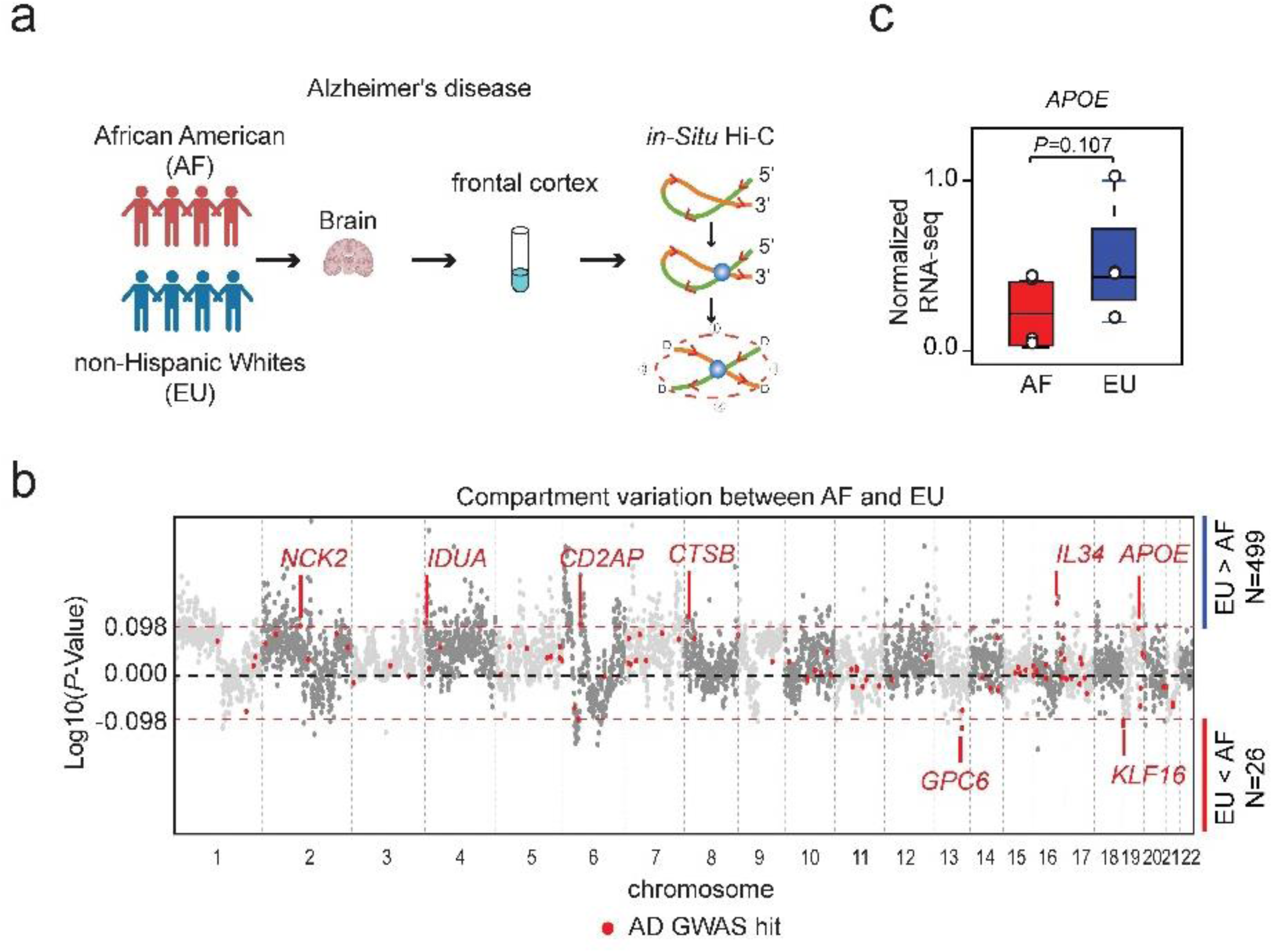
Compartment variations between AF and EU ancestry. **a**, Scheme of experiment design: frontal cortex from four African American and four non-Hispanic Whites donors were subject to in-situ Hi-C analysis. **b**, Scatterplot showing compartment variation between AF and EU ancestry at 500-kb resolution. Each grey dot is a 500-kb bin, while red dots indicate bins containing AD GWAS hits. x-axis: chromosome; y-axis: −log10(P-value), where positive values indicated EU compartments are more active than AF and negative values indicated AF compartments are more active than EU. There are 499 more active compartments in EU and 26 more active compartments in AF ancestry background. **c,** Pseudo-bulk gene expression levels of *APOE* extracted from frontal cortex from the AF and EU ancestry donors used in the Hi-C analysis. Consistent with the compartment analysis, *APOE* has higher expression in frontal cortex from EU ancestry than AF ancestry.

Although the majority of the compartments are relatively consistent across the genome between different ancestries, our observation of more active compartment bins on EU genome than AF genome is consistent with our previously published single nuclei ATAC-seq data from frontal cortex, in which we observed that chromatin accessibility is higher in EU than in AF genome^19^. Using snRNA-seq data from the same brain samples, we previously identified 664 differentially expressed genes across all cell clusters (adjusted *p*-value<0.05;|log_2_Fold Change|>0.32) between EU and AF ancestry (ancestry-DEGs)^20^. Among these ancestry-DEGs, 60 genes reside in differentially compartmentalized bins (**Supplementary Table 2**). For example, *APOE* resides in a compartment bin that is more active in EU than in AF brain tissues which is consistent with previous findings that APOE has significantly greater RNA expression and chromatin accessibility in EU than AF ^19^.

### Variable chromatin loops between AF and EU ancestry

We performed chromatin loop analysis at a 5-kb resolution and evaluated loop variation between AF and EU ancestry. More than 1M loop pixels were called using *DeepLoop* across all eight samples, each pixel representing a pair of 5kb bins that interact with each other. Out of these loop pixels, we identified ∼12K putative EU-specific and ∼3K putative AF-specific loop pixels (*t-test* p<0.05 and > 2-fold change, **Fig. 2a**).

**Figure 2.**
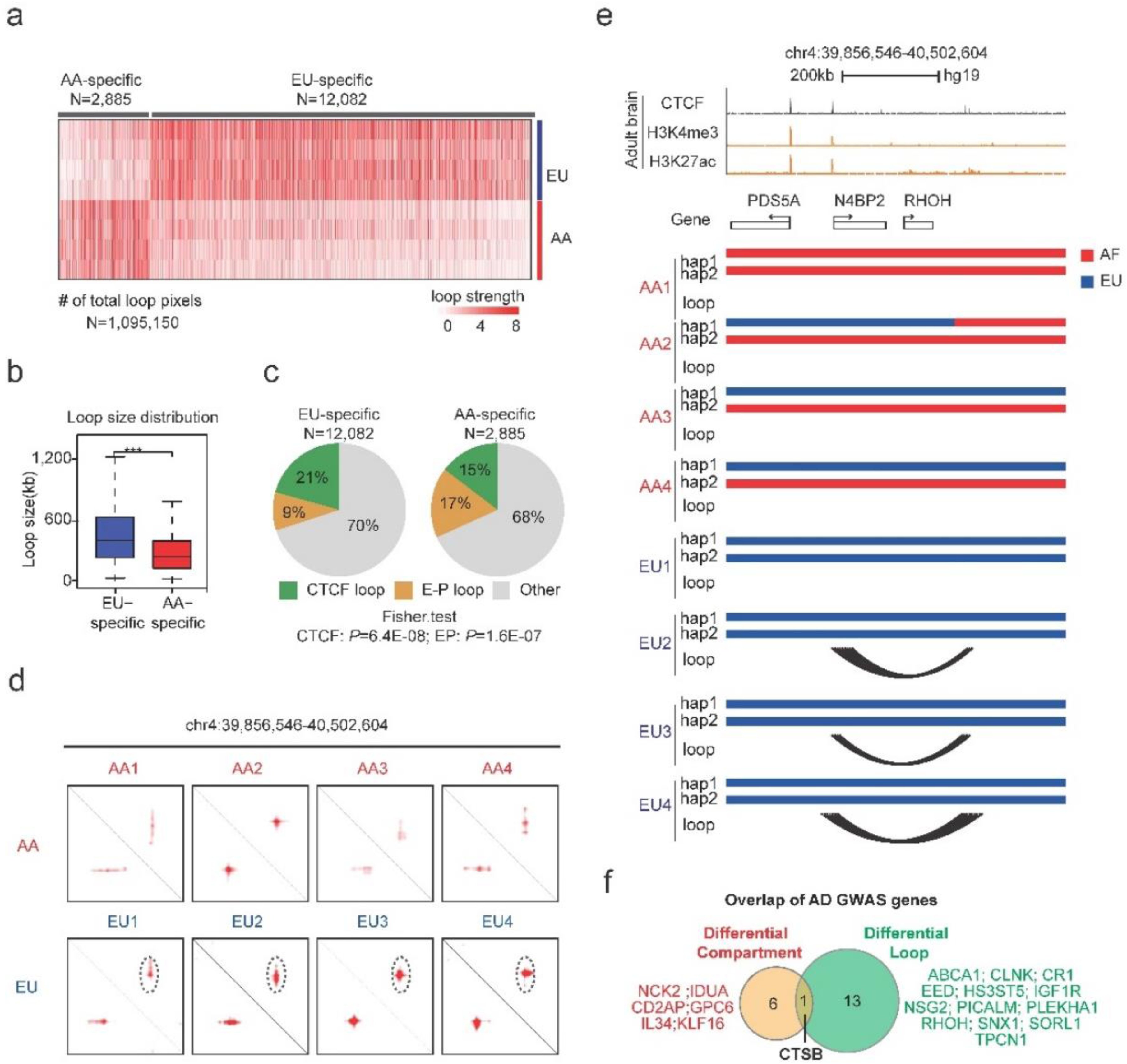
Chromatin loop variations between AF and EU ancestry. **a**, Heatmap showing ancestry-specific loop between AF and EU genomes. Among over one million chromatin loops observed in our dataset, 2885 loops are putative AF-specific and 12,082 loop are putative EU-specific. Column: loop pixel; row: sample; red color gradient: loop strength. **b**, Bar plot showing loop size distribution of putative EU-specific and AF-specific loops. EU-specific loops are larger than AF-specific loops (median size=416 kb vs 261 kb, Wilcox-rank test, P<0.0001). **c**, Pie chart showing loop category distributions for EU- and AF-specific loops. Putative EU-specific loops have a greater proportion of CTCF loops than the putative AF- specific loops (21% vs 15%), whereas the proportion of EP loops is ∼2 fold greater in putative AF-specific than in putative EU-specific loops (17% vs 9%). **d**/**e**, An example of putative EU- specific loop locus shown as Hi-C heatmap (**d**, circled), and arcs on genome browser tracks (**e**). top: CTCF, H3K4me3 and H3K27ac ChIP-seq data in adult brain ^7^; hap1& hap2: homologous chromosomes; red horizontal bar: AF ancestry; blue horizontal bar: EU ancestry; arcs: chromatin loop between two loop anchors. **f**, Venn diagram showing the overlap of AD GWAS risk genes locating at ancestry-specific compartments and ancestry-specific loops.

We characterized the size distribution of ancestry-specific loop pixels and found that Putative EU-specific loops are significantly longer than AF-specific loops (median size=416 kb vs 261 kb) (**Fig. 2b**). Further, we examined the loop anchors of the ancestry-specific loops by assessing their colocalization with CTCF binding sites as well as with brain specific H3K27ac and H3K4me3 ChIP-seq peaks, which represent enhancer and promoter marks, respectively. We then characterized the loops as CTCF-loops if both loop anchors co-localized with a CTCF peak; and EP-loops as those that have both loop anchors co-localizing with either H3K27ac or H3K4me3 peaks. This analysis revealed that the putative EU-specific loops have a greater proportion of CTCF loops than the putative AF-specific loops (21% vs 15%), whereas the proportion of EP loops is ∼2 fold greater in putative AF-specific than in putative EU-specific loops (17% vs 9%) (**Fig. 2c**).

The total sum of ancestry-specific loops (14,967) represents 904 loci (each locus often contains multiple loops). These ancestry-specific loci overlap with 159 out of the 664 ancestry-DEG noted by Griswold et al ^20^. As such, ancestry-DEGs are significantly enriched at ancestry-specific loop loci (Fisher-test; p<2.2×10^-16^; OR=5.13). Fourteen AD GWAS loci are located within ancestry-specific loops (**Supplementary Table 3 & Supplementary Fig 1)**. **Fig. 2d** illustrates one of the GWAS risk loci, i.e. *RHOH*, that resides inside a putative EU-specific loop.

### Identify candidate SNPs that may contribute to ancestry-specific chromatin loops

We hypothesized that genetic variation accounts for at least some of the ancestry-specific loops. In order to link the loop variation to genetic variants, we firstly developed a pipeline to call SNP genotypes directly from the Hi-C sequencing data for each individual (**Fig. 3a & Methods**). On average, we called approximately 4.5 ± 0.17 million SNPs from each AA sample and 3.8 ± 0.16 million SNPs from each NHW sample (**Supplementary Table 1**). Genetic similarity assessed using the Jaccard index revealed greater intra-ancestry similarity compared to inter-ancestry comparisons (**Fig. 3b**). Notably, AA samples have greater genetic diversity between individuals than NHW samples. These observations support the validity of our SNP calling approach and the ancestry of the samples.

**Figure 3.**
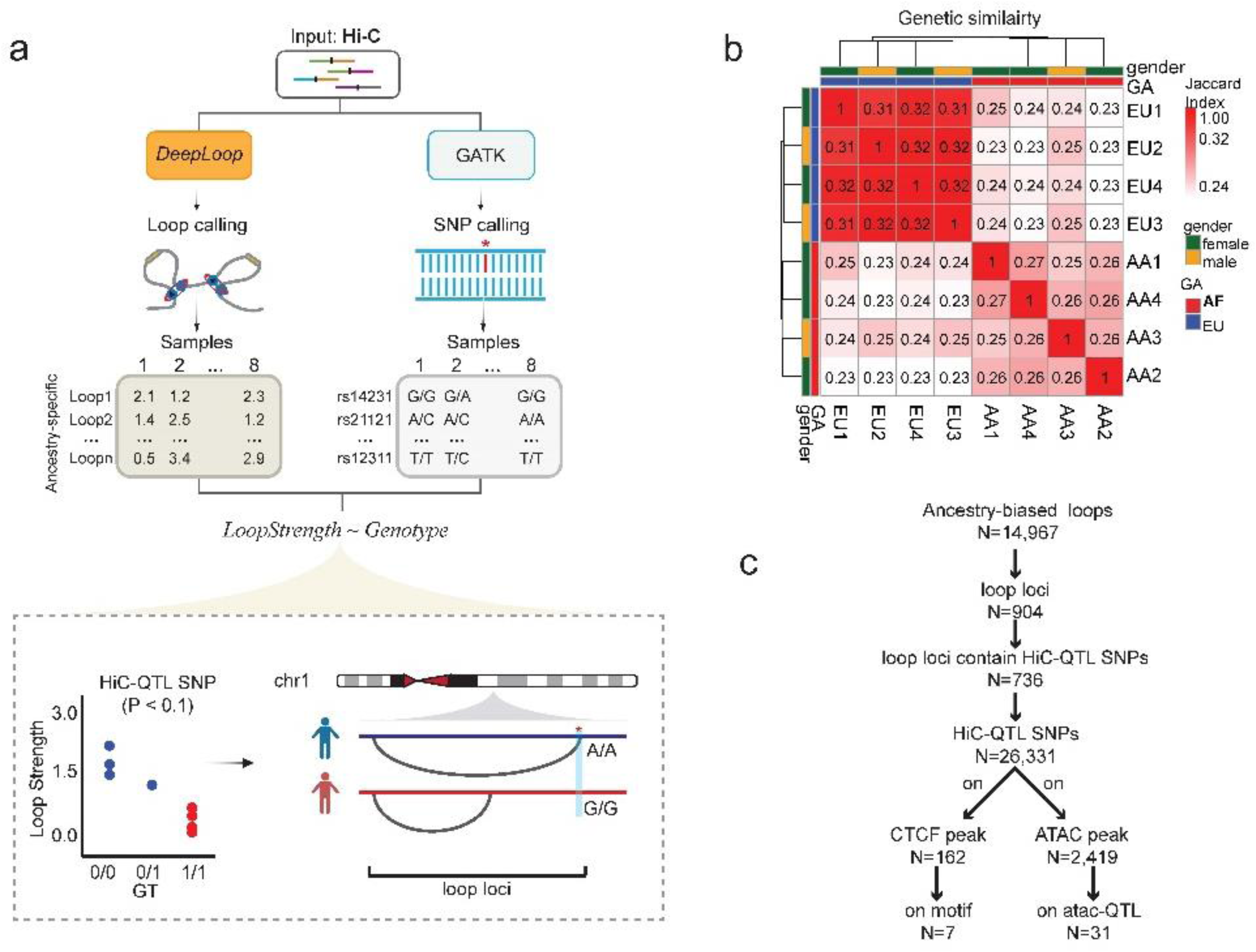
Identification of ancestry HiC-QTLs. **a**, Workflow showing loop calling, SNP calling, and identification of HiC-QTLs. Bulk Hi-C data is used as input for the following steps: 1) perform loop calling using *DeepLoop* and SNP calling using GATK; 2) construct a loop pixel-by-sample loop strength matrix and a variant-by-sample genotype matrix; 3) conduct association analysis between genotypes and loop strengths; 4) output includes significance levels (*P* < 0.1) for associations between each loop pixel and variant. Bottom panel is a carton illustration of a chromatin loop with a HiC-QTL. **b,** Heatmap showing genetics similarity of each donor measured by Jaccard Index. AA samples have greater genetic diversity between individuals than NHW sample. **c**, Characteristics of HiC-QTLs. 1) identification of ancestry-specific loops through pairwise comparisons (total: 14,967 loops); 2) aggregation of loop pixels into loop loci (∼14 pixels per locus); 3) number of SNPs significantly associated with loop pixels; 4) number of SNPs overlapping with CTCF and ATAC peaks; 5) number of SNPs located within CTCF consensus motifs and overlapping with atac-QTLs.

Next, we performed a QTL-like analysis to identify the SNPs associated with the strength of the ancestry-specific loop pixels (**Fig. 3a**). We focused on SNPs residing within the loop anchors based on the hypothesis that a SNP may contribute to differential loop strength by affecting the binding of loop forming factors (CTCF, transcription factors, and cohesion, etc) at loop anchors. Since we expected high false negative rates due to the small sample size, we used a relaxed p-value cutoff (P < 0.1) to be more inclusive than exclusive at this stage. This led to the identification of 26,331 SNPs (HiC-QTL SNPs) (an anchor may have multiple SNPs) that have suggestive evidence for association with 12,550 ancestry-specific loop pixels (HiC-QTL loops), representing 736 out of 904 ancestry-specific loop loci (**Fig. 3c**)^19^.

### Population-specific SNPs may alter Hi-C loops through disrupting CTCF motifs

CTCF has a well-established role in 3D chromatin organization as the blocker of loop extrusion^26, 27^. We integrated epigenomic data from independent ChIP-seq data^7^ for CTCF binding sites to filter our potential HiC-QTLs. We found 162 HiC-QTL SNPs located within CTCF peaks (**Fig. 3c and Supplementary Table 4**). Among these, seven SNPs directly alter the cognate CTCF binding motif, suggesting that these SNPs may alter loop strength via interfering with CTCF binding (CTCF-HiC-QTLs) (**Supplementary Fig. 2)**. **Fig. 4** illustrates an example of a CTCF-HiC-QTL: a long-range loop (>500 kb) is predominantly observed in EU genomes but is largely absent in AF genomes (**Fig 4a, b**). SNP rs2816057 is associated with the strength of this putative EU-specific loop, with the C allele being associated with stronger loop strength than the G allele (**Fig. 4c**). The C allele of rs2816057 is rare in AA population [minor allele frequency (MAF) = 0.017] but common in European populations (MAF=0.41) (**Fig. 4e**). Notably, six out of seven of the identified CTCF-HiC-QTL SNPs (**Supplementary Fig. 2)** show significant differences in MAF between AA and EU populations. In contrast, only 13.5% of genome-wide SNPs show such a MAF difference between populations (p < 0.05, two-sided Fisher’s exact test), suggesting enrichment of population-specific SNPs among CTCF-HiC-QTL SNPs, which supports their role in contributing to ancestry-specific chromatin loops.

**Figure 4.**
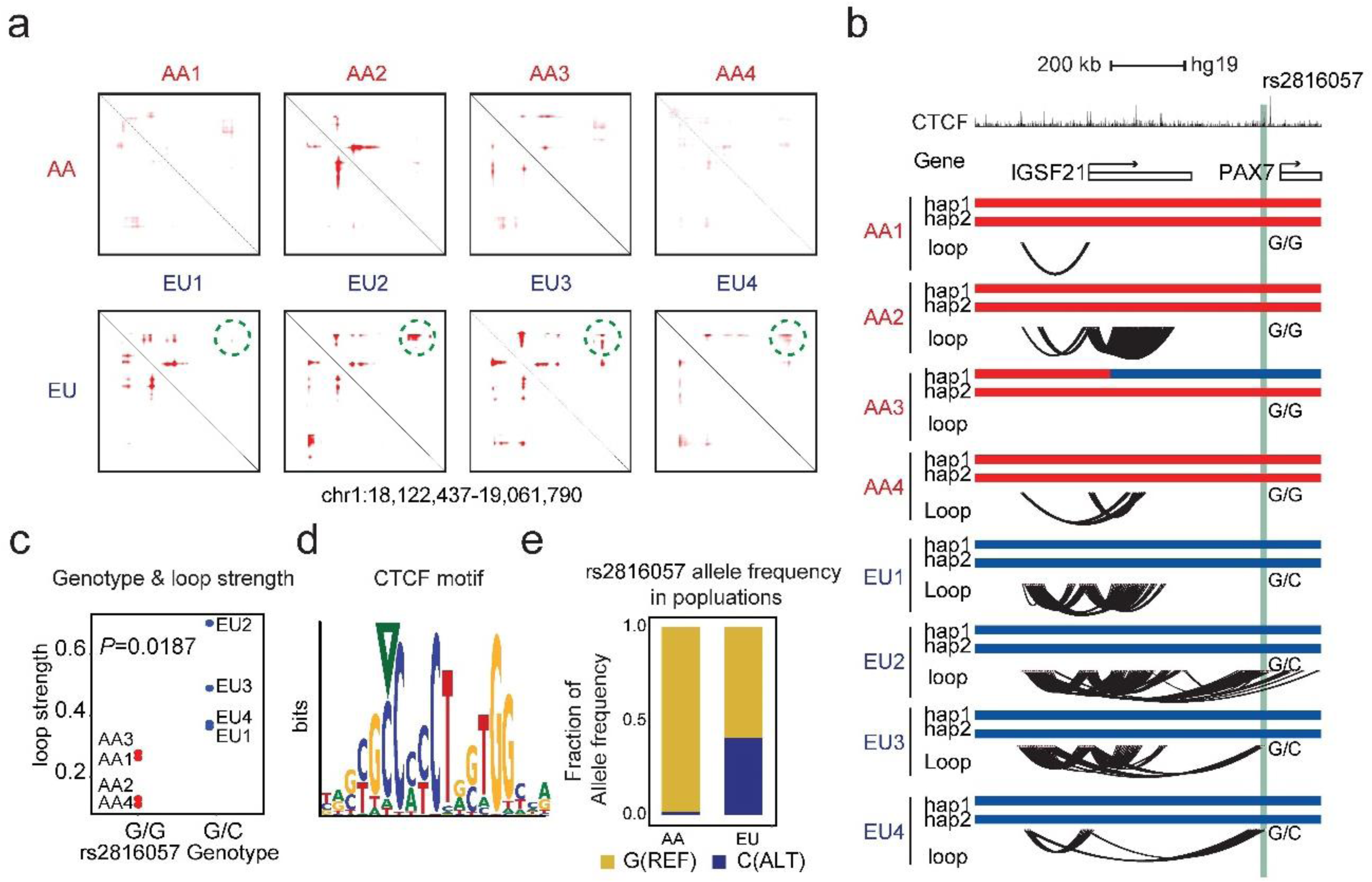
Illustration of CTCF-HiC-QTLs. **a&b,** Heatmap and arc diagram illustrate a putative EU-specific loop (circled in the heatmap) with a CTCF-HiC-QTL (rs2816057, indicated by the green vertical bar in the arc diagram); **c,** Scatterplot showing association between CTCF-HiC-QTL rs2816057 and chromatin loop strength. The C allele of rs2816057 is associated with stronger loop strength; **d,** Position of rs2816057 within the CTCF motif consensus sequence (indicated by the green triangle), where C is the most frequent base and G is almost never observed. **e**, Bar plot showing rs2816057 allele frequency in AA and EU populations. The C allele of rs2816057 is rare in AA population (MAF) = 0.017) but common in European populations (MAF=0.410).

We further hypothesized that at the CTCF-HiC-QTLs, the allele that is conserved in the consensus CTCF motifs should be associated with stronger loop signal. For example, rs2816057 is located at the 6^th^ base of a CTCF motif, where C is the most frequent base while G is almost never observed (**Fig. 4d**). This is consistent with our observation that the rs2816057_C allele is associated with stronger loop signal than the rs2816057_G allele (**Fig. 4c**). To generalize this observation, we extended the analysis to all seven CTCF-HiC-QTL SNPs. In six out of seven CTCF-HiC-QTL SNPs, the allele that disagrees with the canonical CTCF motif (rs2816057-G, rs6862008-A, rs11571379-A, rs12216661-G, rs36007201-C, rs8103622-T, rs73580157_A) is associated with weaker loops than the other allele that is matching the consensus motif sequence (**Supplementary Fig. 2**), indicating a causal effect of disrupted CTCF binding on disrupting Hi-C loops. The p-value for this observation is 0.06 (6 out of 7 trials, binomial test with probability = 0.5). Taking together, our results suggest that population-specific SNPs could contribute to ancestry-specific chromatin loops by altering CTCF binding at loop anchors, and we consider these seven motif-altering SNPs as high-confident CTCF-HiC-QTLs.

### Population-specific SNPs may modulate chromatin loops through effects on open chromatin

Open chromatin regions (OCRs) are enriched with regulatory DNA sequences bound by sequence-specific transcription factors. We reasoned that a HiC-QTL SNP is more likely to have an impact on chromatin loops if it also affects chromatin accessibility. We previously published OCR maps in the same brain samples with ATAC-seq data ^19^. Colocalization analysis found 2,419 putative HiC-QTL SNPs located within brain OCRs, including 31 SNPs that are also associated with variable ATAC peak strength (*i.e.*, these SNPs are also putative ATAC-QTL SNPs) (**Fig. 3c & Supplementary Table 5). Fig. 5** shows an illustrative example of a putative HiC-QTL SNP that is also a putative ATAC-QTL SNP. Rs2001911 resides within the anchor of a putative EU-specific loop (**Fig. 5a&b**) and an ATAC-peak that is more prominent in EU than in AF ancestry (**Fig. 5f**). The G allele is associated with both stronger loop strength and increased ATAC-peak signal (**Fig. 5c&d**). Furthermore, the G-allele is less frequent in the AA population than in the EU population (MAF = 0.13 vs 0.56, **Fig. 5e),** suggesting that it might account for the weaker chromatin loops and ATAC-peak in the AF genome.

**Figure 5.**
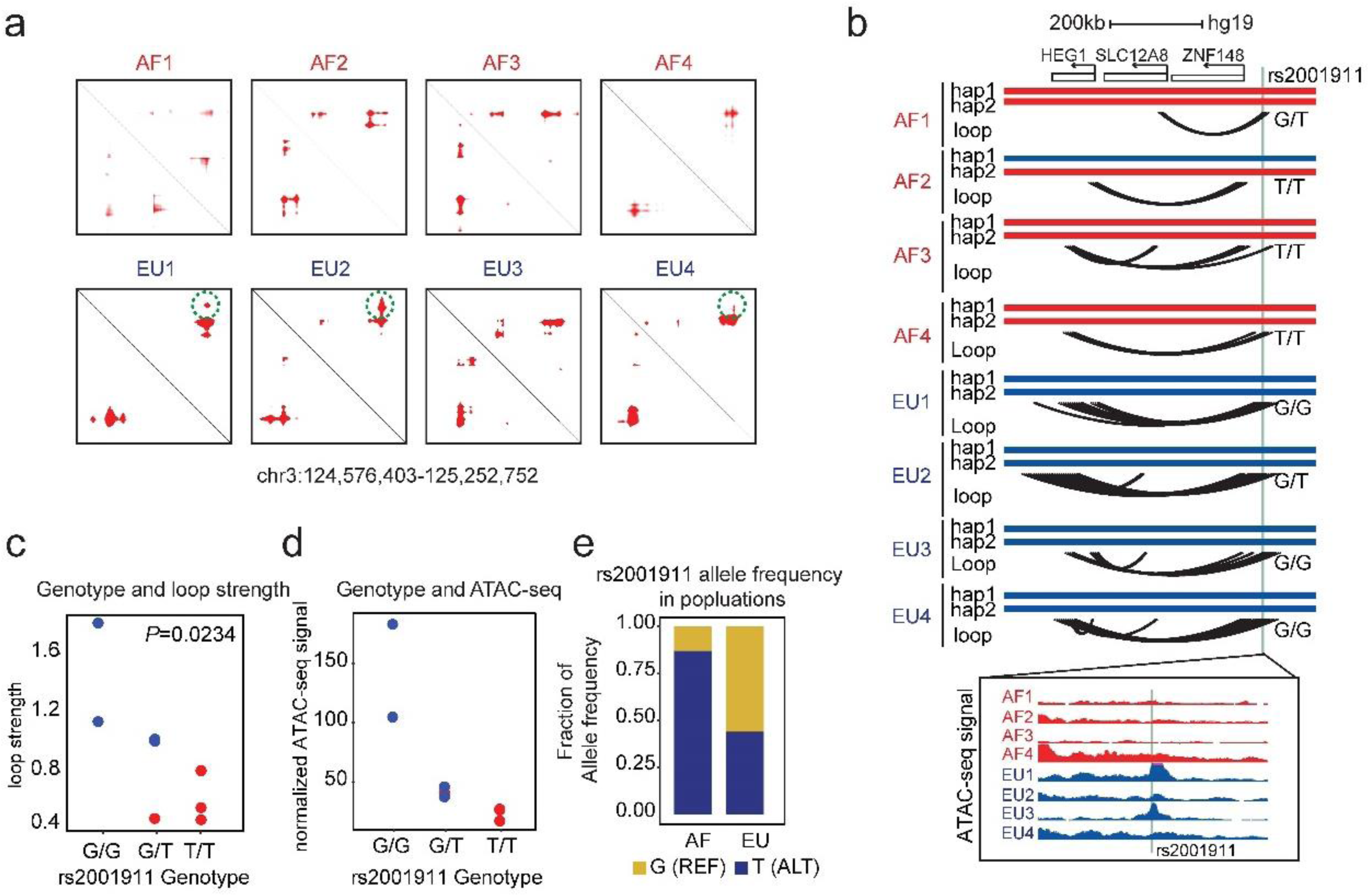
Illustration of ATAC-HiC-QTLs. **a**, Heatmap showing a putative EU-specific loop with an ATAC-HiC_QTL (circled). **b**, Arc diagram illustrating the same loops shown in the heatmap. The Green vertical bar indicates the position of ATAC-HiC-QTL (rs2001911) for an EU-specific loop. Bottom insert displays UCSC track showing ATAC-seq signal around rs2001911 between EU and AF ancestry in frontal cortex; EU genome has higher chromatin accessibility than the AF genome around rs2001911. **c,** Scatterplot showing association between the ATAC-HiC-QTL genotype and chromatin loop strength; G allele of rs2001911 is associated with stronger loop strength. **d,** Scatterplot showing association between the ATAC-HiC-QTL genotype and chromatin accessibility as measured by ATAC-seq; G allele of rs2001911 is associated with higher chromatin accessibility. **e**, Barplot showing rs2001911 allele frequency in AA and EU populations. The G-allele is less frequent in the AA population than in the EU population (MAF = 0.13 vs 0.56).

Among the 31 ATAC-HiC-QTL SNPs, 15 showed different MAF between the AA and EU populations (Fisher-test; p<0.05) and 29 showed concordant effect direction, i.e. the allele associated with stronger ATAC peak is also associated with stronger Hi-C loop (**Supplementary Fig. 3**). This observation strongly suggests a strong link between chromatin accessibility and loop formation (p-value = 1.23 × 10^-7^; 29 out of 31 trials, binomial test with probability = 0.5). We therefore consider the 31 HiC-QTL SNPs as high confidence ATAC-HiC-QTLs that may mediate ancestry-specific chromatin architecture.

## Discussion

Annotating target genes of non-coding GWAS variants remain a challenge as regulatory elements identified by GWAS can be several megabases away from its actual target gene. While expression quantitative trait loci (eQTL) analysis is frequently used to link non-coding polymorphic alleles to changes in gene expression, it is based on statistical association of a specific SNP with a gene expression outcome which could be causal or circumstantial. Most eQTLs capture short-range SNP-gene pairs and it is estimated that only about 20% of GWAS loci have a colocalized eQTL ^36^. 3D genome structure analysis is a valuable complementary tool to the statistical analysis employed in GWAS and eQTL analysis as it detects physical linking of a regulatory element with the promoters of the target gene and is not based on association but on physical interactions ^7^. This spatial information can reveal long-range regulatory relationships hard to detect in linear genome analyses.

Hi-C studies in different cell types, cell states and tissues have revealed dynamic 3D genome architecture during stem cell differentiation, organism development and disease progression, including neurological disorders ^6, 28–30^. Despite the importance of 3D genome structure in maintaining the proper cellular and tissue function, few studies have examined variations of 3D genome structure in the human population as a potential epigenetic mechanism underlying complex disease risk. In this study, we performed an analysis exploring 3D genome structure in postmortem brain tissues as a mechanism of understanding genetic architecture of AD in AF and EU ancestries. Our data showed differences in chromatin interactions between African and European ancestries.

Previously, most studies in 3D genome structure have been done in European ancestry with few studies examining differences in chromatin interactions between AF and EU ancestries However, recent GWAS studies have shown substantial differences in the location as well as variation in the odds ratios for shared associations in African, African American, Peruvian (Amerindian) and European AD populations^10, 13, 21, 31^. Thus, it is increasingly important to understand how ancestry may impact 3D genome structure. Pettie et al examined ancestral differences in CRE in lymphoblastoid cells using the Activity by Contact model, where the chromatin activity and loops were examined using ATAC-seq and H3K27ac HiChIP, respectively ^32–34^. They reported differential chromatin activity between ancestries but did not look for chromatin loop difference between ancestries. Unlike H3K27ac HiChIP that captures chromatin interactions specifically associated with H3K27ac-marked regions, our study directly investigates the ancestry-specific loop interactions unbiasedly using genome-wide Hi-C analysis. This is possible due to several technical improvements: (i) we used *DeepLoop* to improve the quantitation of loop strength; (ii) we further filtered the putative associated SNPs by their ability to impact CTCF motif or ATAC-peak strength. Although our study only identified a small number of high-confidence associations, we expect that a bigger sample size will improve sensitivity.

The increased open chromatin in Europeans vs African genomes seen in the current study supported our previous findings of increased chromatin accessibility in astrocytes of European AD patients versus those of African ancestry using single nuclei ATACseq in frontal cortex^19^. However, ancestry-related differences in chromatin accessibility are not consistently reported across studies. For example, Bhat-Nakshatri et al. examined chromatin accessibility in breast tissue across multiple ancestries and did not find any significant differences, though their analysis is limited to a handful of genes ^35^. While it seems unlikely, further studies would be useful to see if this accessibility difference is unique to neuronal tissue. It is likely that the ancestral difference in chromatin accessibility is due to a complex interaction of factors, including recombination effects (as seen by the smaller haplotypes in Africans relative to Europeans) and the known increased genetic heterogeneity of African ancestries versus Europeans, both which would affect transcription binding sites as well as histone expression and chromatin architecture. Also diet, nutrition, environmental pressures and even the genetic bottleneck that occurred with the migration of individuals from Africa to Europe could also contribute.

To explore the genetic basis of the ancestral-related chromatin loop differences, we conducted a HiC-QTL analysis. To facilitate this analysis, we developed a pipeline to call SNP genotypes directly from the Hi-C sequencing data. Consistent with previous studies^37^, we observed more SNPs in AA than in NHW individuals. We observed that some ancestry-specific chromatin loops are at least partially under genetic control, governed by ancestry-specific SNPs. Identifying QTLs usually requires large datasets, which is currently challenging for Hi-C studies. Given the limited sample size in the current study, we chose to apply an integrative approach to identify high-confidence Hi-C QTLs, where potential Hi-C QTLs were filtered and screened to see if they also were predicted to alter CTCF binding or chromatin accessibility, i.e. CTCF-HiC-QTL SNPs or ATAC-HiC-QTL SNPs

Importantly, we found that many of these high-confidence, regulatory SNPs exhibit significant differences in allele frequency between AA and EU populations—86% of CTCF-HiC-QTLs and nearly half of ATAC-HiC-QTLs show population-specific allele distributions. In contrast, only 13.5% of genome-wide SNPs show such a MAF difference between populations. Moreover, 35 out of 38 high-confidence QTLs (CTCF-HiC-QTLs or ATAC-HiC-QTLs) demonstrated concordant effects on chromatin activity (CTCF binding or accessibility) and loop strength, a pattern highly unlikely to occur by chance. These findings suggest that population-specific SNPs can shape 3D genome architecture in a manner that may influence gene regulation and disease susceptibility.

One notable example is rs34365278-G, a high-confidence HiC-QTL SNP that overlaps with known AD GWAS gene (*CTSB*). Remarkably, we have recently shown that *CTSB* showed differential expression and chromatin accessibility between ancestries, using induced pluripotent stem cell (iPSC)-derived microglia from individuals of AF, EU, and Amerindian ^38^. These findings support the idea that regulatory variation at the *CTSB* locus may contribute to ancestry-specific AD risk, particularly through mechanisms involving chromatin structure and gene expression. This convergence of genetic, epigenomic, and structural variation highlights the potential of 3D genome mapping to refine the functional interpretation of GWAS loci, particularly in underrepresented populations.

Taken together, our work highlights the significance of 3D genome variations in the research of AD genetics and also serves as a proof-of-principle for population-level genetic analysis of chromatin loops with kb-resolution Hi-C analyses.

## Methods

### Brain samples

Brain autopsy samples used in this study have been described previously ^20^. Briefly, all had a confirmed diagnosis of AD upon neuropathological examination and are homozygotes of APOE ε4. Autopsy material was obtained from the Alzheimer’s Disease Research Centers (ADRC) at Emory University, Northwestern University, and the John P. Hussman Institute for Human Genomics (HIHG). All samples were acquired with informed consent for research use and approved by the institutional review board of each center. Global and genome-wide local ancestry (LA) were assessed using genome-wide genotyping data as previously described^15^. Frontal cortex was analyzed from eight donors, four African American and four non-Hispanic White individuals, who were age and sex matched.

### In situ Hi-C library preparation and sequencing

In situ Hi-C library was prepared using a protocol adapted from Rao et al ^25^. Briefly, frozen tissue (≈100 mg) from Brodmann area 9 was pulverized and crosslinked with formaldehyde (final concentration is 1%). The crosslink reaction was quenched with glycine (final concentration of 0.2M). After cell lysis, nuclei were permeabilized with 0.5% SDS and quenched with 1% Triton X-100. DNA was digested with Mbo I (New England Biolab) in situ overnight. After restriction digestion, the DNA ends were filled in and labeled with biotinylated-dATP (Active Motif) before proximity ligation. Crosslinkes were reversed and the ligated DNA purified. DNA was sheared using Covaris LE220 (Covaris, Woburn, MA). DNA fragments in the range of 300-500 bp were enriched with AmPure XP beads (Beckman Coulter). Ligation junctions with biotin label were pulled down with streptavidin beads (Invitrogen) and prepped for Illumina sequencing with 8 cycles of PCR amplification. For each library, 340∼860 million of paired-end reads at 150 bp length were obtained.

### Hi-C data processing

Hi-C sequencing reads were trimmed to 36 bp. R1 and R2 raw reads were mapped to the hg19 reference genome separately using bowtie^39^ (v.1.1.2). Two output SAM files were then paired together with an in-house analysis script. After removal of PCR duplicates, we discarded reads with both ends mapped to the same *DpnII* fragments. All remaining valid read pairs represented two different *DpnII* fragments. Because cut-and-ligation events are expected to generate reads within 500 bp upstream of *DpnII* cutting sites due to size selection (‘+’ strand reads should be within 500 bp upstream of a *DpnII* site and ‘−’ strand reads should be within 500 bp downstream of a *DpnII* site), we retained only read pairs with both ends satisfying these criteria. We next split all remaining valid read pairs into three classes based on their strand orientation (‘same-strand’, ‘inward’ or ‘outward’). We retained inward read pairs if the distance between two ends was <1 kb, and outward read pairs if the distance between two ends was <5 kb. We further merged these filtered inward, outward and same strand as the *cis* reads pair. The *HiCorr* ‘*DpnII*’ mode was used to acquire bias-corrected 5-kb anchor loop files from *cis* and *trans* fragment read pairs.

### Compartment analysis

We first performed a lower-resolution (500 kb) compartment-level analysis partitioning the genome into compartment A (associated with transcriptionally active euchromatin) and compartment B (associated with transcriptionally silenced heterochromatin) following the method described previously ^4^. The “state” of compartmentalization was measured with the first principal component (PC1) values^25^. To identify differences in the 3D genome structure between EU and AF ancestry at the compartment level, we performed pair-wise t-tests. Firstly, we constructed contact matrix at 500-kb resolution for each chromosome. Secondly, we normalized the matrix by using the genome distance then generated a correlation matrix based on normalized value for each chromosome. Further, the principal component analysis on the correlation matrix then assigns the genome into two compartments by using PC1. Then, we use the density of transcription start sites (TSS) of each 500-kb bin to determine compartment A and compartment B.

First, we calculated the average PC1 value for the four African (AF) patients and the four European (EU) patients separately. Next, we computed the difference between the average PC1 values of the AF and EU patients, defining this value as Diff_compartment. We then utilized a simple Z-score to identified significant different region (with *P*<0.05; *cutoff*=0.015; two-side).

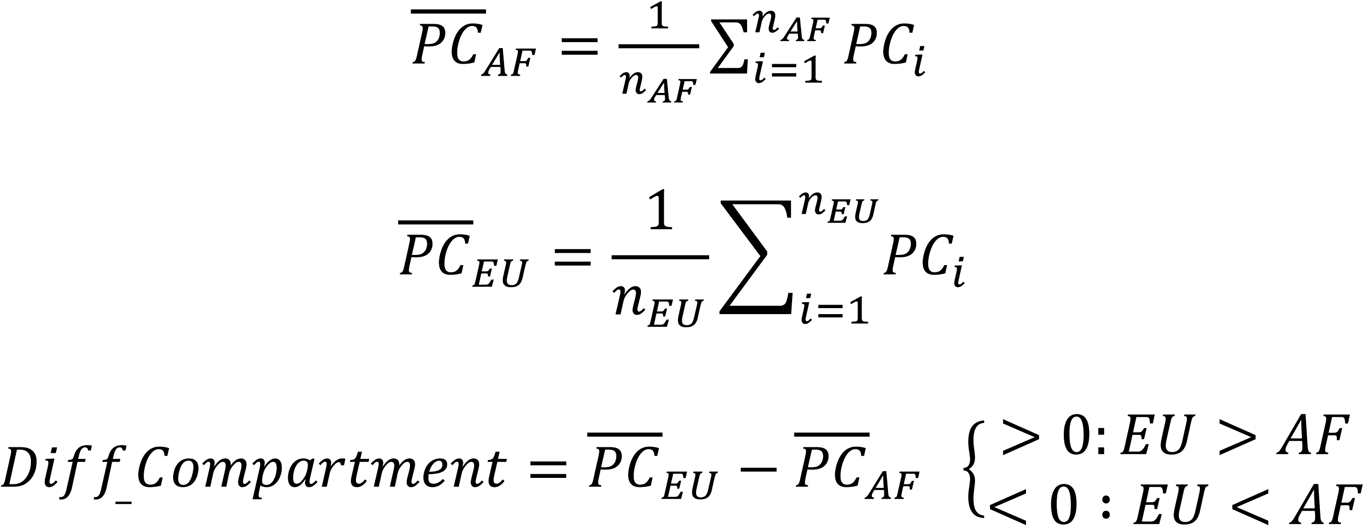

### Chromatin loop analysis

Unbalanced sequencing depth of Hi-C datasets can introduce biases for downstream quantitative comparisons across individuals. To mitigate this issue, we took the sample with lowest depth (cis-contacts within 2M) as a baseline to down-sample the rest of all higher-depth Hi-C data. These depth-balanced matrices were then used for subsequent analyses, including *HiCorr bias correction*, *LoopEnhance* as well as loop comparison across two populations.

Firstly, we utilized *HiCorr* pipeline to perform both explicit and implicit bias correction. Unlike normalization methods, which preserve a strong diagonal signal in the contact heatmaps, *HiCorr* corrects distance effects in a joint function with other biases and outputs the observed/expected ratio heatmaps for chromatin interaction profiling. Secondly, we chose a proper *LoopEnhance* model based on sequence depth of cis-contact within 2M and then achieved a robust chromatin loop calling.

After *HiCorr* bias correction and *LoopEnhance*, (1) we took top300k loops from each patient and then generate a union matrix *M*_*ij*_ (i is row number and each row refers to one loop pixel; j is column number and each column is one sample) and each element of matrix is loop strength of a given loop pixel for a given patient; (2) we performed supervised two-side T-test to identify significant differential loop pixel between AF and EU AD patients (*P < 0.05; Fold-change > 2*).

For each loop pixel, it was linked by two different 5-kb anchors referring to two different genomic loci. Loop size is measured by the relative distance between these two different genomic loci.

### Definition of different types of chromatin loop

By using supervised approach, we obtained EUR-specific and AFR-specific loops. To distinguish what types loop they are, we integrated these loops with ChIP-seq data, mainly CTCF and Enhancer/Promoter. (1) we intersected loop anchor with CTCF and Enhancer/Promoter peak; (2) Loop pixel with two anchors overlapping with CTCF peaks is called as CTCF loop; (3) Loop pixel with at least one anchor overlapping with enhancer and promoter is called as E-P loop.

### Haplotype and SNP calling

Local ancestry at each locus across the genome was calculated using RFMix with SNP-array data. A reference panel with diverse populations from 1000 Genomes was used for to assign European (EU), African (AF) and Amerindian (AI) ancestries.

All pair-end Hi-C data in this paper is aligned to hg19 (human reference genome) by using bwa with default parameters. Non-uniquely mapped reads have been removed by using samtools^40^. PCR-duplications derived from library construction are depleted by using *Picard.* INDEL is called by using GATK-HaplotypeCaller with default parameter. Full SNP information table is generated by using VCFtools.

Jaccard index

The genotypes of any two samples are taken from a set of eight samples. The Jaccard Index is then calculated to measure the similarity between these two samples. This is done by dividing the size of the intersection of their genotypes by the size of the union of their genotypes.

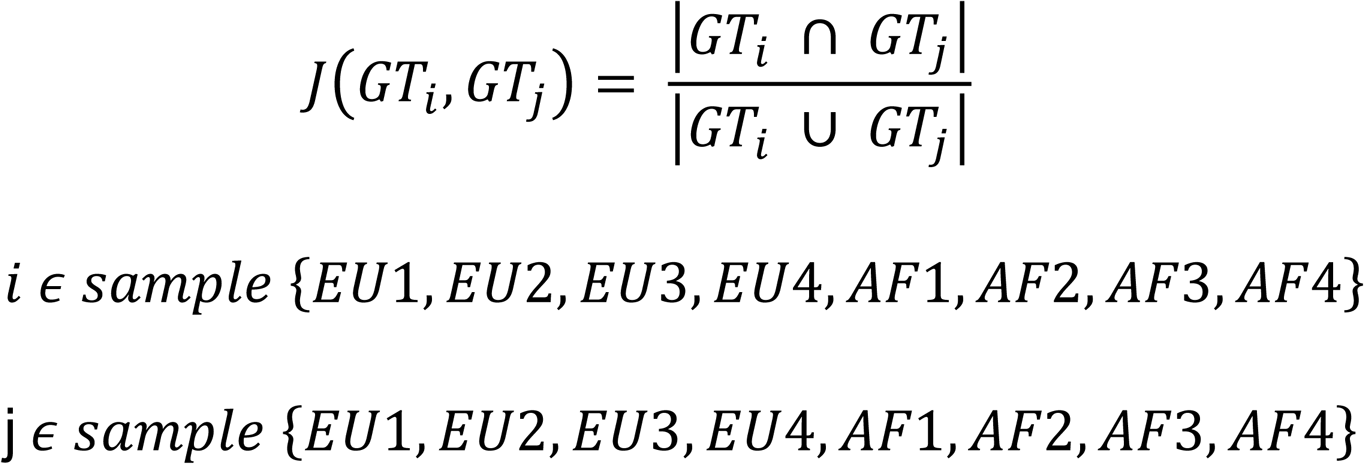

### HiC-QTL SNPs analysis

From loop analysis, we generated one matrix Mij (i is row number and each row is one loop; j is column number, and each column is one sample; each element of matrix is loop strength). From SNP analysis, we generated full SNP table Nij (i is row number and each row is one SNP; j is column number, and each column is one sample; each element of matrix is genotype information). To link SNP to loop, we first generated a reference file to link SNP to 5-kb anchor to obtain SNP-to-anchor reference file. And then, link this reference file with chromatin loop, of which each loop pixel has two anchors. To quantify the association between genotype and loop, linear regression analysis is used as below:

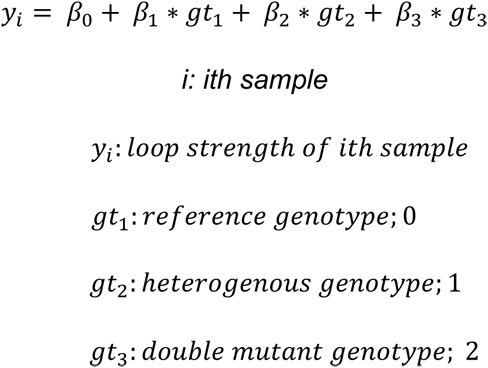

### ChIP-seq

Adult brain ChIP-seq data are public data published in a previous study ^7^. Firstly, all sequencing data was aligned to hg19 by using bowtie. After the mapping step, we then remove non-uniquely mapped and PCR duplication read from the original bam file. Next, we took clean bam file and then use Macs2 to call peak. Finally, we generate bigwig files to achieve data visualization on UCSC track.

### ATAC-QTL SNPs analysis

Our ATAC-seq data identified 226,237 OCRs in the genome and 169,642 OCRs have at least one SNP. Out of the 514,067 SNPs within these OCRs, from pseudo-bulk ATACs-seq analysis, we firstly called ATAC peak of each sample and then created one union peak matrix M where each row represents one ATAC peak and each column represents one AD brain sample; each element of matrix is normalized ATAC signal). From SNP analysis, we generated full SNP table Nij (i is row number and each row is one SNP; j is column number, and each column is one sample; each element of matrix is genotype information). To link SNP to ATAC peak, we first generated a reference file to link SNP to each peak to obtain SNP-to-peak reference file. And then, link this reference file with each atac peak. To quantify the association between genotype and atac peak signal, linear regression analysis is used as below:

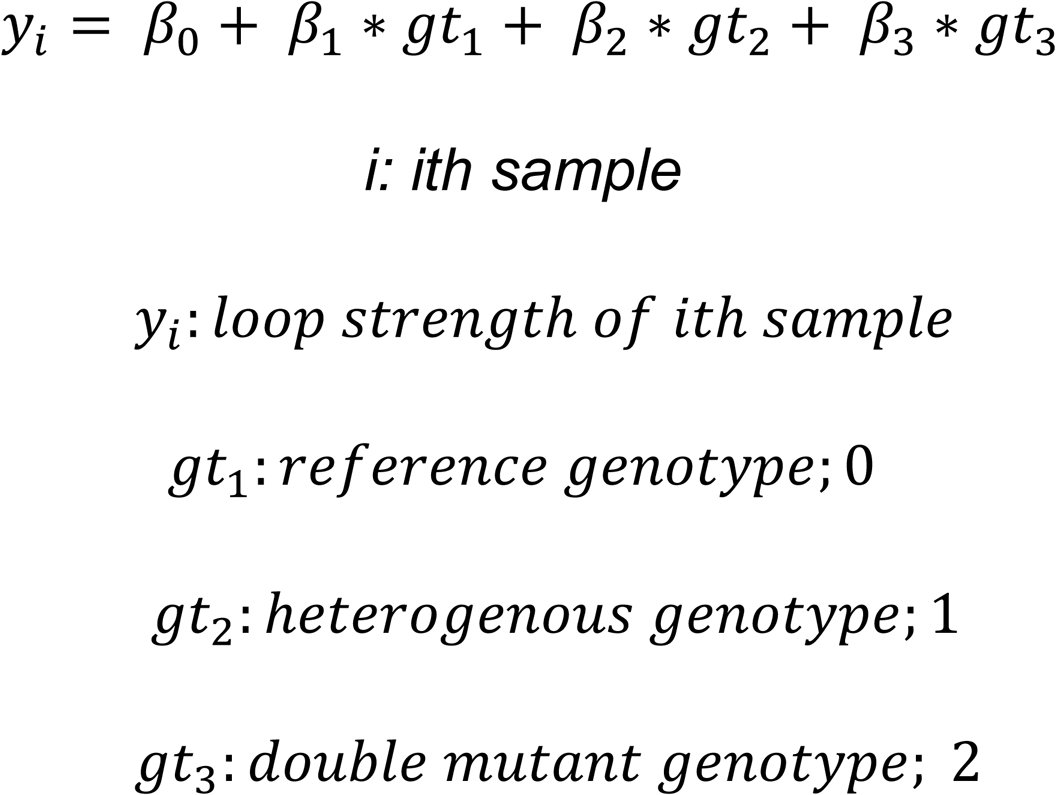

### Definition of loop loci

We define the location of each loop pixel as the entire region between the left and right anchors (including the anchors themselves). To define ancestry-specific loop (AB-loop) loci, we merge all AF-specific and Putative EU-specific loop pixels as we identified using the “merge” option in bedtools; each AB-loop locus usually has multiple loop pixels. Loop pixels with gap size < 20kb will be merge into one locus.

### Population-specific HiC-QTL SNPs

We downloaded the summary table of SNPs from dbSNP database with allele frequency among diverse populations. We searched our Hi-C-QTL SNPs within this table and then calculated the odd ratio of a given SNP between European and African American. In this case, we would be able to confirm whether Hi-C-QTL SNPs display population specificity.

### CTCF motif calling

Adult brain CTCF peak and canonical CTCF binding motif (downloaded from Hocomoco) as input files are under-processed by fimo. In this result, we obtain adult brain CTCF binding motif and its direction.

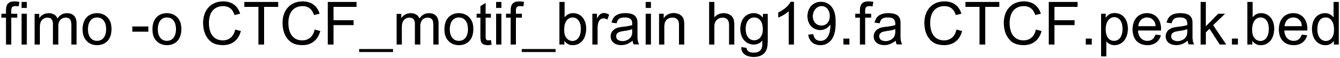

## Acknowledgments

This study was supported by the National Institute of Aging (grant numbers R01-AG070864, U01-AG072579, RF1-AGO59018) and National Human Genome Research Institute (R01-HG009658, R01-HG012384).

We acknowledge the Center for Genome Technology (CGT) from the John P. Hussman Institute for Human Genomics (HIHG) from the University of Miami, Miller School of Medicine for the genomic and data analyses. We express our gratitude to the participants, researchers, and staff involved for their invaluable contributions to the present study.

---

**Supplementary Figure 2.**
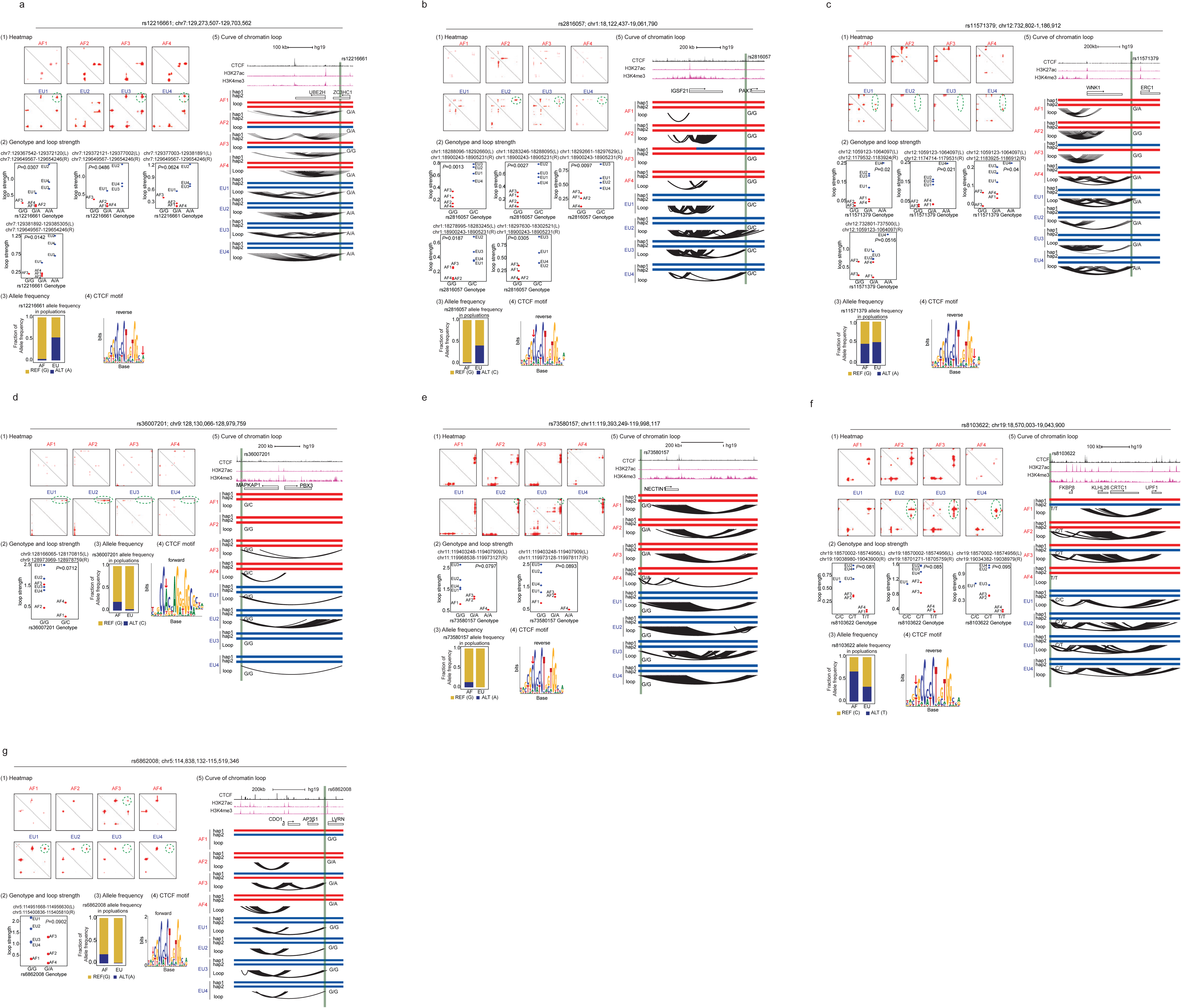
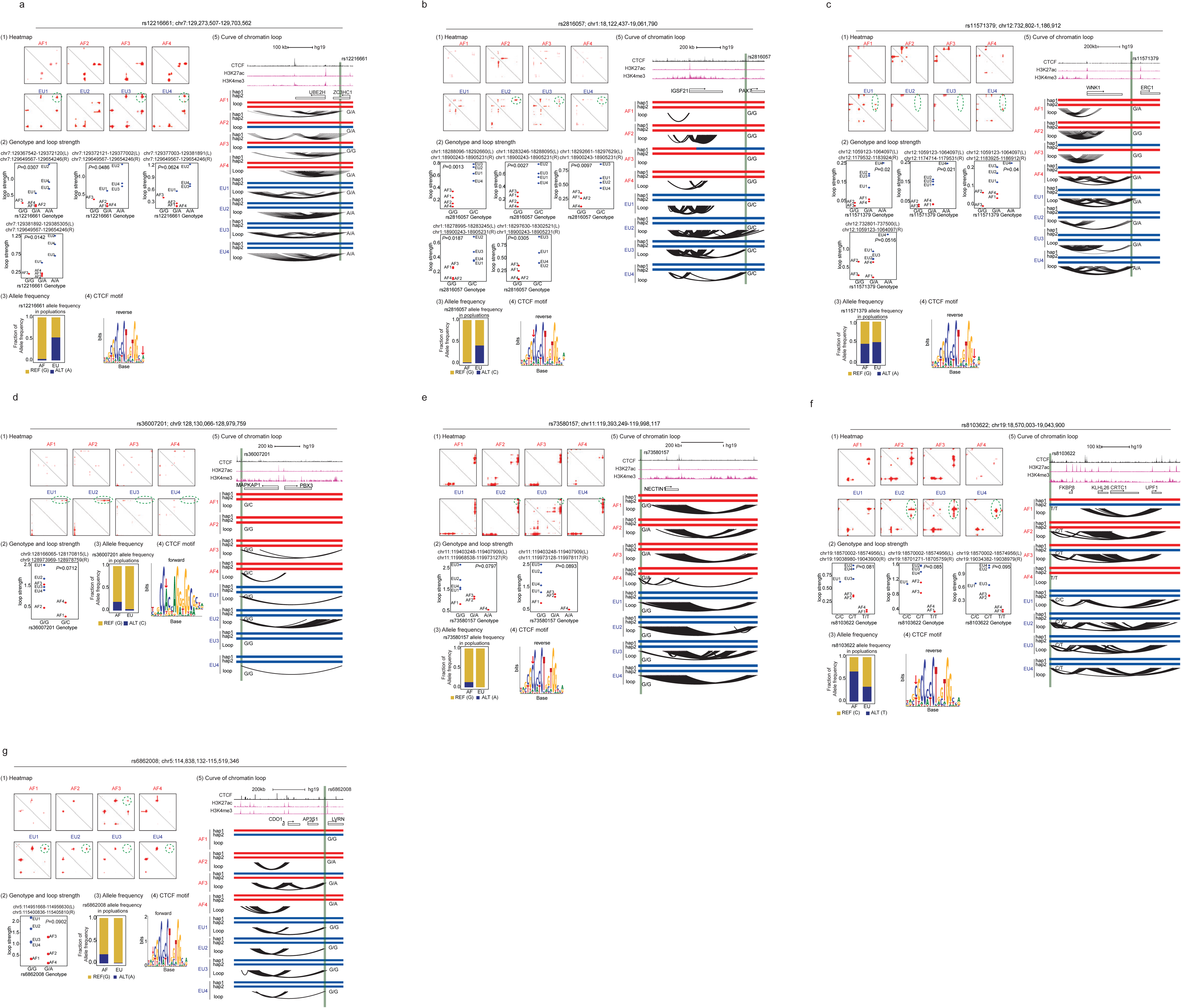
Illustration of CTCF-HiC-QTLs

**Supplementary Figure 3.**
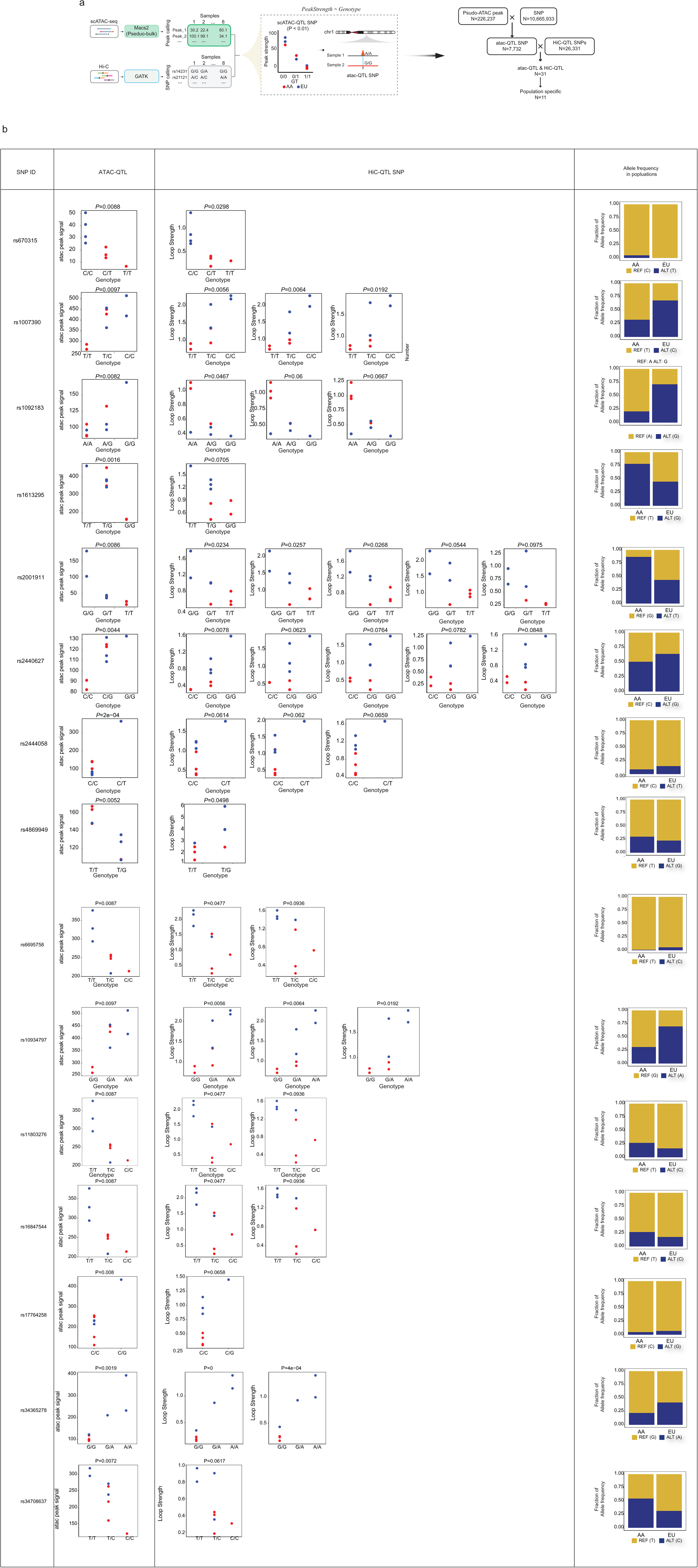
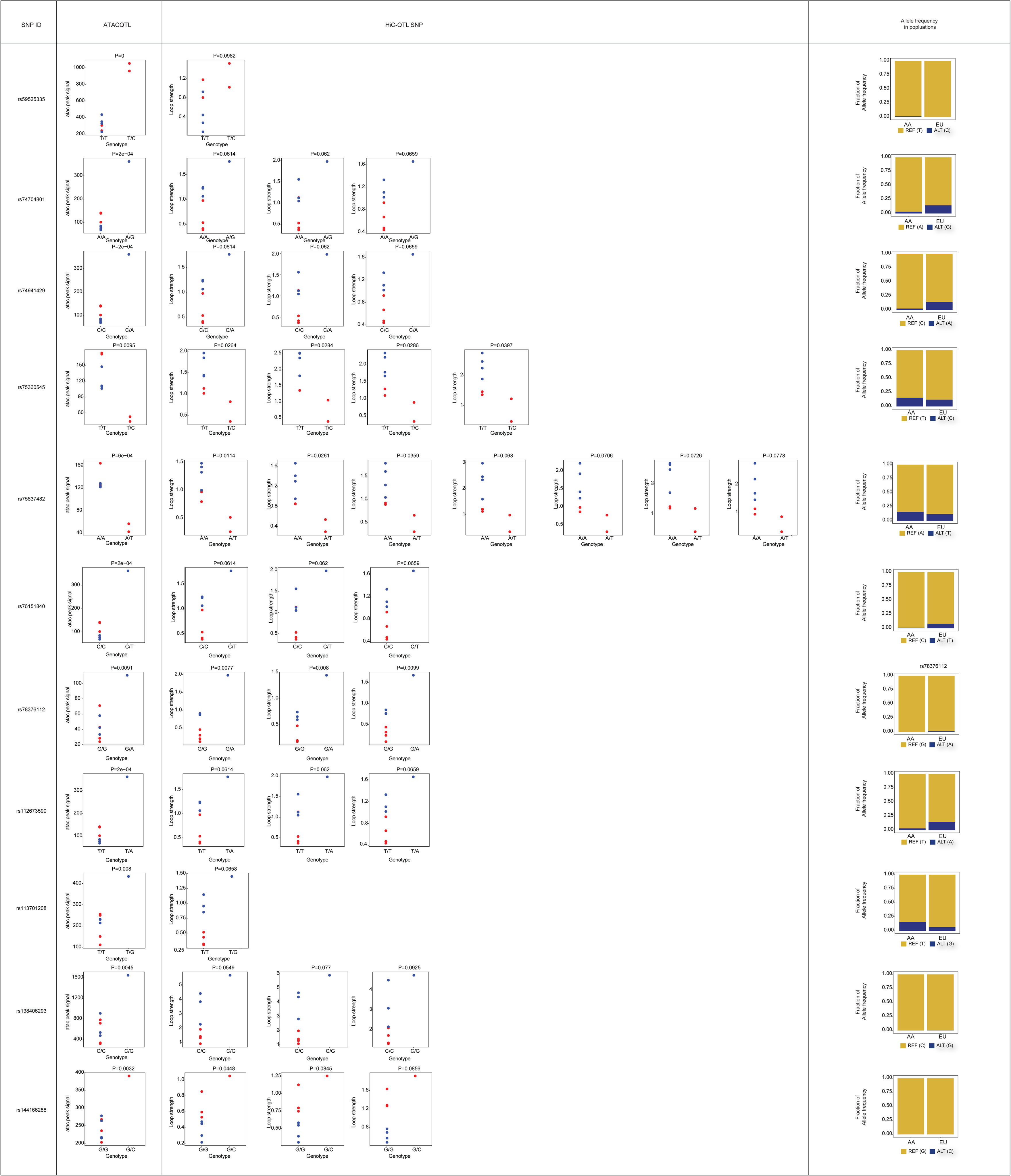
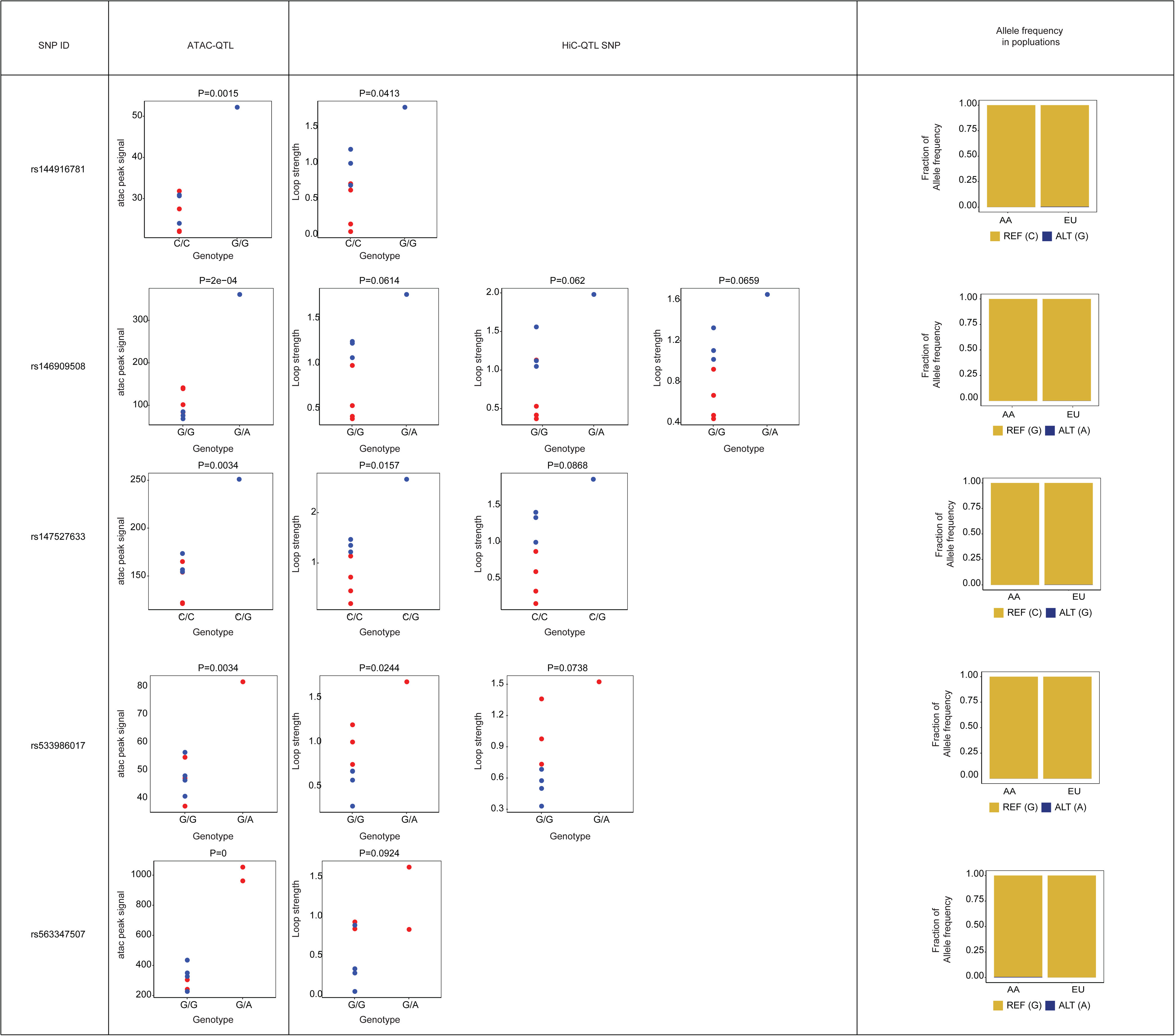
Illustration of ATAC-HiC-QTLs

